# GIVE Statistic for Goodness of Fit in Instrumental Variables Models with Application to COVID Data

**DOI:** 10.1101/2021.04.18.440376

**Authors:** Subhra Sankar Dhar, Shalabh

## Abstract

Since COVID-19 outbreak, scientists have been interested to know whether there is any impact of the Bacillus Calmette-Guerin (BCG) vaccine against COVID-19 mortality or not. It becomes more relevant as a large population in the world may have latent tuberculosis infection (LTBI), for which a person may not have active tuberculosis but persistent immune responses stimulated by Mycobacterium tuberculosis antigens, and that means, both LTBI and BCG generate immunity against COVID-19. In order to understand the relationship between LTBI and COVID-19 mortality, this article proposes a measure of goodness of fit, viz., <u>G</u>oodness of <u>I</u>nstrumental <u>V</u>ariable <u>E</u>stimates (GIVE) statistic, of a model obtained by Instrumental Variables estimation. The GIVE helps in finding the appropriate choice of instruments, which provides a better fitted model. In the course of study, the large sample properties of the GIVE statistic are investigated. As indicated before, the COVID-19 data is analysed using the GIVE statistic, and moreover, simulation studies are also conducted to show the usefulness of the GIVE statistic.

## 1 Introduction

There has been a significant interest since COVID-19 outbreak on whether well known tuberculosis (TB) vaccine Bacillus Calmette-Guerin (BCG) and COVID-19 mortality are related or not. Strictly speaking, the scientists are willing to know the effect of BCG vaccination on the reduced mortality from COVID-19. Though as of now, there is no as such any scientific evidence in favouring the positive impact of BCG vaccination on the mortality due to COVID-19 (see, e.g., Soliman et al. (2020)). However, most of the studies did not consider the fact that many people in this world may have latent TB infection (LTBI), that means, those people may not have evidence of clinically active TB but have Mycobacterium tuberculosis antigens. Moreover, the fact is that LTBI creates lifelong immunity, and hence, may provide an immunological protection against COVID-19. This article addresses the association between LTBI and the reduction of mortality due to COVID-19, and in the course of this study, the Instrumental Variables (IV) estimation has played an important role in the modelling of COVID-19 data to draw such conclusions, see Takahashi (2020). However, the conclusions drawn in that article are restricted only to the estimation of parameters, and there is no mention/evidence of goodness of fit, i.e., how good the model is fitted to the COVID-19 data. In this context, we would like to mention that this article attempts to propose and study the goodness of fit of the models obtained by IV estimation, which has not yet been paid much attention in the literature.

We would now like to discuss in detail about how IV can be a powerful toolkit to work on the aforesaid research problem related to COVID-19 outbreak. Note that one may formulate a regression equation with the mortality due to COVID-19 as the response variable and a few related covariates, which will be thoroughly discussed in Section 5. However, in such regression models, it is likely to happen that the covariates become correlated with the regression errors, and it makes the regression model a dubious regression model. In order to overcome this problem, one may adopt the IV method since the IVs are highly correlated with the covariates but uncorrelated with the error random variables, at least in the limiting sense. An excellent exposition about the IV estimation is presented in Bowden and Turkington (1984) and Wansbeek and Meijer (2000, Chap. 6).

Apart from the COVID-19 outbreak, in general also, the IV estimation in multiple linear regression models has played a vital role in biostatistics. Bio-statisticians are pleased to use it as IV can tame the confounding bias, without directly observing all explanatory variables. Baiocchi (2014) has presented a tutorial on IV estimation in biostatistics. The IV estimation has not only been used but extended in many directions. For example, Martinussen and Vansteelandt (2020) used IV estimation in competing risk data, Tchetgen et al. (2015) extended it to censored survival outcome, Li and Brookhart (2015) applied IV techniques to the proportional hazards model, Dai (2014) presented a test for checking concordance between instrumental variable effects, Tchetgen (2014) presented a method of instrumental variable estimation when the outcome is rare, and many others (see also Burgess et al. (2017), Didelez (2007), Uddin et al. (2015)).

As we indicated earlier in the context of the modelling mortality rate due to COVID-19 outbreak, the IV estimation in multiple linear regression model requires a set of IVs, which are highly correlated with the explanatory variables, at least in limit and uncorrelated with the disturbance term, at least in limit. Various sets of different IVs can be obtained for the same set of data, which gives rise to different estimators of the regression coefficient vector. This, in turn, produces different fitted models for the same set of given data. A more complicated situation arises in multiple linear regression where there are more than one explanatory variables. In such a case, the experimenter has various options to choose the IVs for the explanatory variables, e.g., the same IVs for all the explanatory variables or different IVs for different explanatory variables. Now, how to check which of the model is better fitted to the given data or equivalently which choice of instrumental variable gives a better fit in statistical sense is an issue. Many occasions, researchers and applied workers provide various arguments to justify their choice of IVs but such justifications are based on their experience and not based on any quantitative and scientific outcome of statistical measures.

It is important to note that the goodness of fit in the classical multiple linear regression model, based on OLSE, is defined for judging the goodness of fit of estimation, denoted as *R*^2^ whereas for the purpose of prediction, it is denoted as 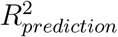. We now would like to emphasize that the goodness of fit for judging the estimation and prediction of the model are different. Strictly speaking, this article considers the aspect of goodness of fit of estimation only while Pesaran and Smith (1994) mentioned that it is problematic to develop the goodness of fit based on residuals, and for that reason, they considered the prediction errors to develop the goodness of fit. We here successfully develop the goodness of fit based on residuals, see also Windmeijer (1995), Bloom, Bond and Van Reenen (2007) in this regard.

Overall, this article investigates in the following direction. The coefficient of determination, popularly known as R-square (*R*^2^), is used to measure the goodness of fit in classical multiple linear regression model and cannot be used or directly extended to the case of IV models because the explanatory variable becomes stochastic and correlated with the disturbance term. To overcome this problem, we here propose a new goodness of fit statistic, viz., <u>G</u>oodness of <u>I</u>nstrumental <u>V</u>ariable <u>E</u>stimates (GIVE) statistic, which can be used to measure the goodness of fit in the IV models and to determine the optimal choice of IVs for a given data. We here consider the development of GIVE statistics based on IV estimation for a general situation when the error random variables are non-spherical, and their covariance matrix is unknown. The statistical properties such as the asymptotic distributions of the GIVE statistic is also derived. Moreover, The performance of the GIVE statistic in a finite sample is investigated with the Monte-Carlo simulation experiments in the context of multiple linear regression model and measurement error model. Finally, the usefulness of the GIVE statistic is established on the COVID-19 data addressing the association between LTBI and the reduction of mortality due to COVID-19 as we discussed in the first two paragraphs in this section.

## 2 Instrumental Variable (IV) Estimation

We first describe the model and IV estimation in the setup of a classical multiple linear regression model, see Rao et al. (2007, Chapter 4). Consider the multiple linear regression model with the study variable *y* linearly related to the *p* explanatory variables with an intercept term in *X* as

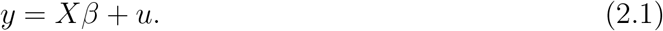

Here *y* is the (*n*×1) vector of observations on study variable, *X* is the (*n*×(*p*+1)) matrix of *n* observations on each of the *p* explanatory variables and an intercept term, *β* is the ((*p*+1)×1) vector of regression coefficients associated with the (*p* + 1) explanatory variables, and *u* is the (*n* × 1) vector of nonspherical disturbances with *E*(*u*) = 0 and unknown covariance matrix, i.e., *E*(*uu*′) = *σ*^2^Ω^−1^, where Ω is an unknown positive definite matrix, and *σ >* 0 is a constant.

Note that in the usual linear regression model, it is assumed that *X* and *u* are uncorrelated but we here consider a situation where *X* and *u* are uncorrelated in limit, in the sense that 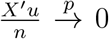 as *n* → ∞, where 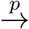 denotes the convergence in probability. Moreover, we assume the presence of intercept term in the model, which is needed for the validity of coefficient of determination in classical multiple linear regression model and may be needed for the validity of the proposed goodness of fit statistic studied in Section 3. Next, suppose that a set of *p* instrumental variables (denoted by (*Z*_1_, . . ., *Z*_*p*_)) is available, and the observations are arranged in a (*n* × (*p* + 1)) matrix with an intercept term in *Z* = (*Z*_0_*, Z*_1_, . . ., *Z*_*p*_) such that they are correlated with *X*, in limit and uncorrelated with *u*, in limit. Assume that

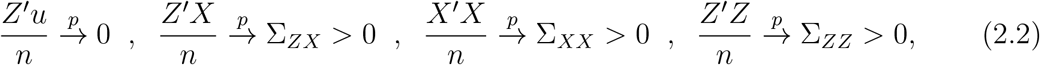

where Σ_*ZX*_, Σ_*XX*_ and Σ_*ZZ*_ are non-singular positive definite matrices of constants, and *I* is an (*n* × *n*) identity matrix.

Consider now the set up of linear regression model in which *X* is regressed on the set of instruments *Z* for a given sample size *n* in the first stage as

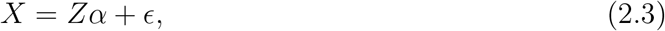

and we assume *E*(*ϵ*) = 0 and *E*(*ϵϵ*′) = *σ*^2^Ω^−1^ for the random error term *ϵ*, where *σ >* 0 and Ω are the same as the earlier associated with *E*(*uu*′). In this context, it should be mentioned that *E*(*uu*′) and *E*(*ϵϵ*′) can be different in principle but here they are considered to be the same only because of the notational and algebraic simplicities. The feasible generalized least squares estimate of *α* is obtained by using generalized regression of *X* on *Z* and replacing Ω by 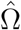 as

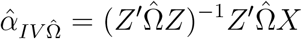

from (2.3), where 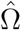 is a consistent estimΩator of unknown Ω. In the next stage, using the the predicted value of *X* as

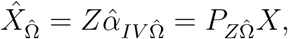

where 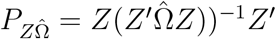 in 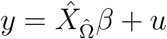, we have

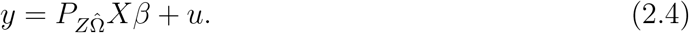

Application of generalized least squares on (2.4) yields the two stage feasible generalized least squares (2SFGLS) estimator of *β* as

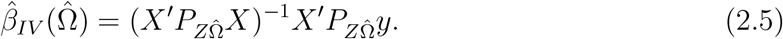

Now, Theorem 1 asserts the consistency of 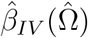 to estimate the unknown parameter *β*. Suppose that 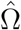 be a consistent estimator Ω, and the following assumptions are required for the consistency of the estimator 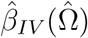:

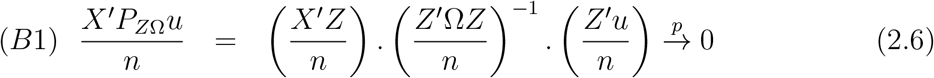

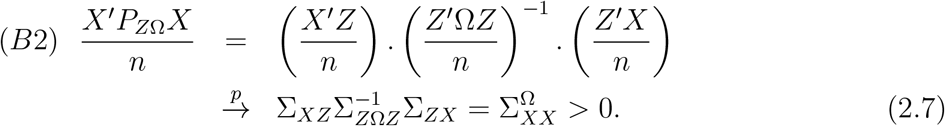

### Theorem 1

*Let* 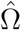 *be a consistent estimator of* Ω *under Euclidean norm. Then, under* (*B*1) *and* 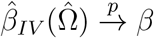 *as n* → ∞.

**Proof:** See the supplementary file.

## 3 GIVE Statistic for Goodness of Fit in IV Model

Note that the coefficient of determination, popularly known as *R*^2^, is the ratio of the sum of squares due to regression (or the fitted model) and the total sum of squares obtained in the context of analysis of variance in multiple linear regression model, see Rao et al. (2008, Chap. 3). It is measuring the proportion of variability explained by the fitted model with respect to the total variability in the data based on the ordinary least squares estimator (OLSE). The coefficient of determination fails to judge the goodness of fit in the IV models because the total sum of squares can no longer be partitioned into two orthogonal components, viz., the sum of squares due to regression and the sum of squares due to error. Moreover, the IV estimators do not possess the properties like the best linear unbiased estimator as possessed by the OLSE, and hence, replacing OLSE by IV estimator in the definition of goodness of fit statistics will not necessarily yield a good and reliable outcome. We here attempt to provide a new statistic for quantitatively measuring the goodness of fit in the IV models, termed as GIVE statistics.

The square of the population multiple correlation coefficient between *y* and fixed explanatory variable, say *X*\* is given by

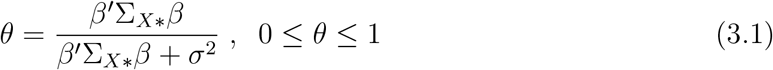

where 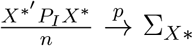, where 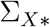 is a positive definite finite matrix, 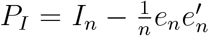, and *e*_*n*_ = (1, 1, . . ., 1)′ is a (*n* × 1) vector of all elements unity. Note that when the model is best fitted, then *σ*^2^ = 0, and we have *θ* = 1. On the other hand, if the model is worst fitted, then all the *β*’s will be zero indicating that all the explanatory variables are not effective, and consequently, *θ* = 0. Any other value of *θ* lying between zero and one will accordingly measure the goodness of the fitted model in terms of multiple correlation coefficient.

Now, recall the model (2.4), viz., 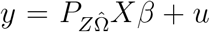, where *β* is estimated by (2.5), and we develop the goodness of fit statistic. The total sum of squares for model (2.4) is

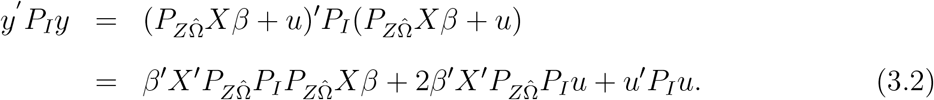

It may be observed that in the classical multiple linear regression model, the total sum of squares is partitioned into two orthogonal components- sum of squares due to regression and sum of squares due to error. Observing the expression in (3.2), the first two terms, viz., 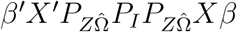 and 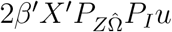 can be considered as jointly constituting the sum of squares due to regression whereas *u*′*P*_*I*_*u* can be considered as the sum of squares due to errors. If we replace the unknown 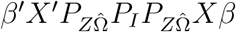 and 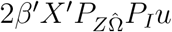 by 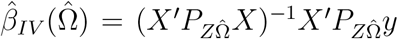 and 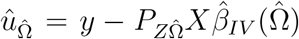, respectively, then a measure of goodness of fit can be constructed by measuring the ratio of the sum of squares due to regression and the total sum of squares under IV models as

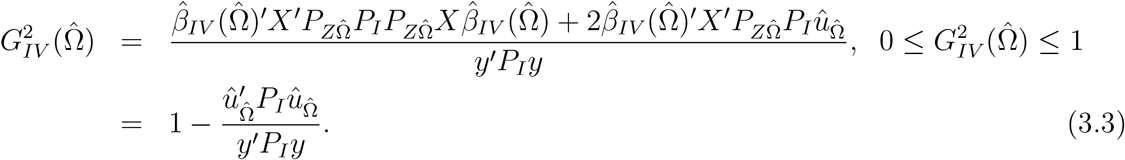

The statistic (3.3) can be used to measure the goodness of fit in the IV model having spherical disturbances with unknown covariance matrix and is termed as <u>G</u>oodness of <u>I</u>nstrumental <u>V</u>ariable <u>E</u>stimates (GIVE) statistic in IV models. Theorem 2 states the consistency of 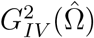 to its population counterpart *θ*_*IV*_(Ω), where

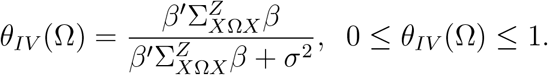

The interpretation of *θ*_*IV*_(Ω) is the same as *θ* in (3.1).

### Theorem 2

*Let* 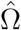 *be a consistent estimator of* Ω *under Euclidean norm. Then under* (*B*1) *and* (*B*2), 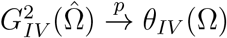 *as n* → ∞.

**Proof:** See the supplementary material.

**Remark:** In case Ω is known, then 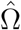 can be replaced by known Ω. For example, substituting Ω = *σ*^2^*I* will give the classical case.

In all-inclusive, 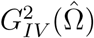 is a consistent estimator of *θ*_*IV*_(Ω), where unknown Ω is estimated by its consistent estimator of Ω. The interpretations of 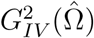 are similar to the interpretations of coefficient of determination. When *σ*^2^ = 0, then 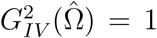 indicates that the model is the best fitted. On the other hand, if all estimated regression coefficients are close to zero or say, exactly zero which indicates that all the regression coefficients are not significant, i.e., the model is worst fitted, then 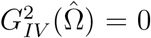. Beside these two values of 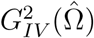, any other value of 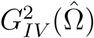 lying between 0 and 1 will indicate the degree of goodness of fit provided by the fitted model for given explanatory variables and sample size. For example, if 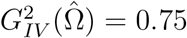, it then would indicate that 75% of the variation in the response values is being explained by the fitted IV model based on the choice of IVs and the fitted IV model is nearly 75% good.

### 3.1 Asymptotic Distribution of GIVE statistic

We here derive the asymptotic distribution for the case when Ω is unknown, and the asymptotic distribution for the special cases, viz. known Ω and Ω = *σ*^2^*I* are stated in Corollary 1 and Lemma 1 (see the supplementary file), respectively.

We now assume the following conditions for the sake of technicalities:

(A1) The parameter space of *β* is compact.

(A2) *X* is a bounded random variable.

(A3) *Z* is a bounded random variable.

(A4) *Z* and *u* are independent random variables.

(A5) *Z* and *E* are independent random variables.

(A6) The correlation between *ϵ* and *u* is non-zero.

(A7) Let 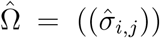 and Ω = ((*σ*_*i,j*_)), where 1 ≤ *i, j* ≤ *p*. Moreover, suppose that 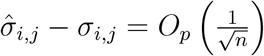 for all *i* and *j*.

Before stating the theorem, we also need to introduce a few notations, which are the following. Let us denote *a* = (*a*_1_, . . ., *a*_*d*_) ∈ ℝ^*d*^, which is an arbitrary *d*-dimensional vector, and

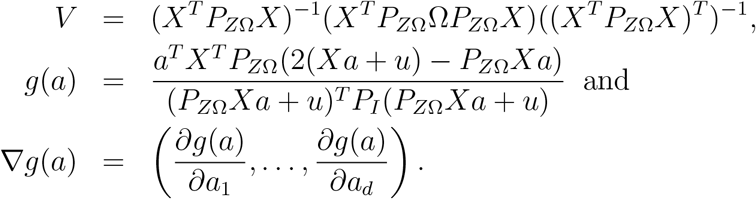

#### Theorem 3

*Under conditions (A1)-(A7)*, 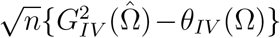 *converges weakly to a normal distribution with mean* = 0 *and* 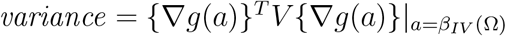.

**Proof:** See the supplementary material.

#### Corollary 1

*Under conditions (A1)-(A6),* 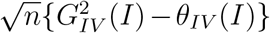 *converges weakly to a normal distribution with mean* = 0 *and* 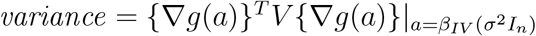.

**Proof of Corollary 1**: The proof follows using the same arguments as the proof of Lemma 1 in the supplementary file.

## 4 Monte-Carlo Simulation Study

We conducted a Monte-Carlo simulation experiment for studying the performance of the GIVE statistic for finite sample cases. To justify and understand the behaviour of GIVE statistics based on the choice of instrumental variables, we consider Wald’s and Durbin’s choices of instrumental variables, (see Rao et al. (2008, Chap. 4, pp. 208-209) for more details). The Wald instrument technique divides the observations on explanatory variable into two groups based on their median value and choose the IV as +1 and −1 for the two groups, and Durbin instrument technique uses the ranks of observations on explanatory variable as instruments. Here our modest aim is to understand the performance of the proposed goodness of fit statistics. Hence to simplify and make the understanding better, we consider a model with homoskedastic error structure with identity covariance matrix. The same procedure can be extended to the case of a non-identity type covariance matrix-known or unknown both. In case of unknown non-identity covariance matrices, we also need to choose a suitable consistent estimator to estimate the covariance matrix based on a sample of data.

We conducted the simulation experiments using R software under two cases. First case is the general set up of multiple linear regression models. In this case, our objective is to demonstrate the application of GIVE statistics in a prominent model where IV is extensively used. Keeping this in mind, we consider the measurement error models in the second case and simulated the values of proposed GIVE statistics.

### 4.1 Application : Multiple Linear Regression Model

Here our simulation set up is as follows. For a given sample size *n* = 50, 100 and 200, the random errors *ϵ* are generated following *N*(0, *σ*^2^*I*_*n*_) with *σ*^2^ = 1.5, 2.5, 3.5, 4.5 and 5 for the model (2.3) and (2.4) with *p* = 5 and 9 with high values of population multiple correlation between the study variable and all the independent variables, and Ω is known. The observations on *X* are generated from a normal distribution and corresponding IV’s are found to construct *Z* using the Wald instrument technique, denoted as *Z*_1_, and Durbin instrument technique, denoted as *Z*_2_. The GIVE statistics based on *Z*_1_ and *Z*_2_ are computed and denoted as 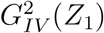 and 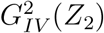, respectively and their empirical relative bias and empirical relative mean squared error are computed. A comparison of the GIVE statistic with traditional *R*^2^ is made to know what happens to GIVE statistic when some important and unimportant explanatory variables are added or when the intercept term is absent etc. It may be recalled that the classical coefficient of determination (*R*^2^) has a property that it increases as the number of explanatory variables in the model increases. The traditional *R*^2^ is defined only when there is an intercept term in the model. *The detailed results along with obtained values in different tables are presented in the supplementary material for the sake of space.* The conclusions drawn from those results are mentioned here.

We observe that as the value of variance *σ*^2^ increases, the values of 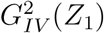 and 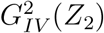 decrease. All the results indicating that the choice of *Z*_2_ yields a better fitted model than *Z*_1_. Such an outcome is intuitively correct also because *Z*_2_ is using more information in terms of ranks of the observations whereas *Z*_1_ is using the information on data as indicator variables only. Hence, it can be concluded that if both *Z*_1_ and *Z*_2_ are used in *X*, then the proposed GIVE statistics are capable of judging the goodness of fit which is affected primarily due to the choice of IVs and thus deciding over the appropriateness of the choice of IVs. We have used the same choice of IVs for all the variables but extending it to a case where different explanatory variables are replaced by different IVs is not difficult. The resulting GIVE statistics will reflect the goodness of fit appropriately.

Moreover, the empirical relative bias (RB) and empirical relative mean squared error (RM) of 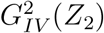 is smaller than that of 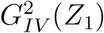 in all the settings of simulation set up. The RB and RM of 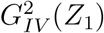 and 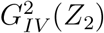 increase as *σ*^2^ increases for the given sample size. As the sample size increases, the RB and RM of 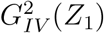 and 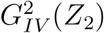 decrease for a given *σ*^2^. It is clear that the behavior resembles the behaviour of traditional *R*^2^. The 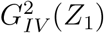 and 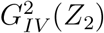 are found to have a tendency to increase as the the number of explanatory variables increase. This empirically confirms that the values of GIVE statistics increase when explanatory variables are added in the model. Next, when relevant explanatory variables are added, both 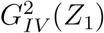 and 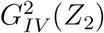 increase. The magnitude of increment is less than the magnitude of increment of 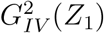 and 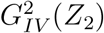. This clearly indicates that the capability of GIVE statistic in diagnosing whether the relevant or unimportant explanatory variables are added in the model.

Furthermore, we find that the values of the GIVE statistic increase when the intercept term is absent in the model in comparison to the values when the intercept term is present in the model. There seems no issue that the GIVE statistics do not work in a model without intercept. It is contrary to the traditional *R*^2^ which is defined only in a model with intercept term. The RBs and RMs of 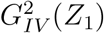 and 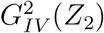 are lowered when the intercept term is removed from the model. It will give slightly higher values in comparison when the intercept term is present in the model.

Besides, the GIVE statistic works well only when *σ*^2^ is small. It is intuitively expected that as *σ*^2^ increases, the model fitting should be worsened. The GIVE statistics are capturing it better than *R*^2^. Difference in the values of 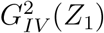 and 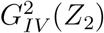 for the same sets of explanatory variables can be interpreted as the difference arising due to the choice of IVs.

### 4.2 Application : Measurement Error Model

We first briefly describe the set up of measurement error models. More details can be found in Rao et al. (2008), Cheng and Van Ness (1999). Fuller (1987). Note that the symbols and notations used in this subsection are limited to this subsection 4.2 only. The reason to choose the measurement error model for application of GIVE statistics is that the explanatory variable and random errors become correlated when the data is contaminated with the measurement errors. Beside, among other estimation methods used in measurement error models, the IV method is a popular method to obtain the consistent estimators of the regression coefficient.

Note that a basic common assumption of any statistical analysis is that all the observations are correctly observed. However, in many practical situations, they cannot be correctly observed due to various reasons; in fact, they are observed with some measurement error into them. The difference between the observed and true values of the variable is termed as measurement error. We here consider the structural form of the multiple measurement error model where the true explanatory variables are stochastic with the same mean. Let *η* = (*η*_1_*, η*_2_*, . . ., η*_*n*_) denote the (*n* × 1) vector of observations on the true values of study variable and *T* = (*t*_1_*, t*_2_*, . . ., t*_*n*_)′, *t*_*i*_ = (*t*_*i*1_*, t*_*i*2_*, . . ., t*_*ip*_)′; *i* = 1, 2, . . ., *n* be the *n* × *p* matrix of the *n* observations on each of the *p* explanatory variables linked with

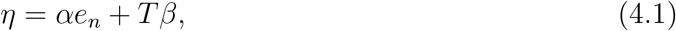

where *β* = (*β*_1_*, β*_2_, . . ., *β*_*p*_)′ is a (*p* × 1) vector of regression coefficients associated with *p* explanatory variables, *α* is the intercept term, and *e*_*n*_ = (1, 1*, . . .,* 1)′ is a (*n* × 1) vector of elements unity. The true values *η* and *T* are not observable due to presence of measurement errors but they are observed as *y* and *X* which are *n* × 1 vector and *n* × *p* matrix, respectively given as

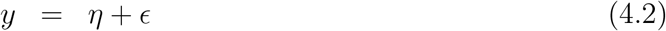

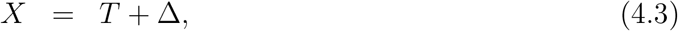

where *ϵ* = (*ϵ*_1_*, ϵ*_2_*, . . ., ϵ*_*n*_)′) is a (*n*× 1) vector of measurement errors in *η*, Δ = (*δ*_1_*, δ*_2_*, . . ., δ*_*n*_)′, and *δ*_*i*_ = (*δ;*_*i*1_*, δ*_*i*2_*, . . ., δ*_*ip*_)′; *i* = 1, 2*, . . ., n* is *n* × *p* matrix of measurement errors involved in *T*. We assume that *δ*_*ij*_ (*i* = 1, 2*, . . ., n*, *j* = 1, 2*, . . ., p*) are independent and identically distributed random variables following 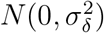, and *ϵ*_*i*_, (*i* = 1, 2*, . . ., n*) are also independent and identically distributed following 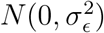. Further, *ϵ* and Δ are also assumed to be statistically independent of each other.

The data is generated from a population with high multiple correlation, and *n* = 50, 100 and 200 are chosen. The measurement errors *ϵ* and *δ* are generated following 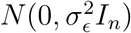 and 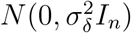, respectively with 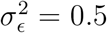, 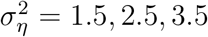 and 5 for the model (4.1)-(4.3) when *p* = 5 and 9. The observations on *T* are generated from a normal distribution in every replication, and the corresponding IV’s are found to construct *Z* using the two approaches - Wald Instrument Technique, denoted as *Z*_1_, and Durbin Instrument Technique, denoted as *Z*_2_. The *β* is estimated using *Z*_1_ and *Z*_2_ with measurement error ridden data on *X* and *y* to further compute the GIVE statistics. In this section, the GIVE statistics based on *Z*_1_ and *Z*_2_ are denoted as 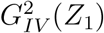 and 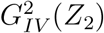, respectively. The results are summarized in the supplementary material.

It is clear that the measurement errors in the data affect the values of 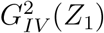 and 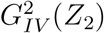. As the value of variance 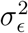 increases, the values of 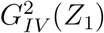 and 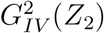 decrease. The rate of such increment depends upon the sample size also. This again confirms that the proposed GIVE statistics can satisfactorily measure the goodness of fit in the measurement error models. The performance of GIVE statistics depends on a combination of *n* and 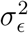. Overall, it can be concluded that if there are several available choices of IVs, e.g., *Z*_1_ and *Z*_2_ in the present case, then the proposed GIVE statistic helps in judging the appropriate choice of IVs and gives an idea about the goodness of the fitted model.

Next, in terms of bias, it empirically indicates that 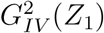 and 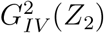 are negatively biased. The magnitude of relative bias of 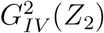 is smaller than the magnitude of relative bias of 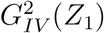 and 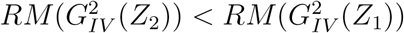 in all the settings of simulation set up. If we take the sample size to be substantially large, then as the sample size increases, the values of GIVE statistics will converge better towards *θ*. Moreover, we observe that the values of 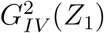 and 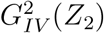 have a tendency to increase as the number of explanatory variables increases just like the traditional *R*^2^ in the classical multiple linear regression model without measurement errors. When unimportant explanatory variables are added in the model, then 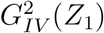 and 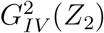, both increase but the amount of increment is less than the increment that happened in 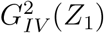 and 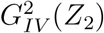. This confirm the capability of 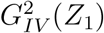 and 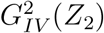 in diagnosing whether the relevant or unimportant explanatory variables are added in the model.

Hence, the proposed GIVE statistics works well in the measurement error model. We expect that the GIVE statistics will also work well in other models also.

## 5 Analysis : COVID-19 Data Set

In Section 1, it has already been mentioned that there is likely to be an association between well known tuberculosis (TB) vaccine Bacillus Calmette-Guerin (BCG) and COVID-19 mortality. However, the scientists have not yet found any data analytic evidence of such a finding (see, e.g., Soliman et al. (2020)). One possible reason for not finding any evidence of it is that a large proportion of people in this universe may have latent TB infection (LTBI), i.e., they may not have clinical evidence of active TB but have Mycobacterium tuberculosis antigens. Moreover, as said before, the LTBI may provide immunological protection against COVID-19. We here try to address the association between LTBI and the reduction of mortality due to COVID-19 using the IV technique based on GIVE statistic.

The data set consists of mortality rate due to COVID-19 of 104 countries from January 01, 2020 to May 30, 2020 (see https://www.worldometers.info/coronavirus) along with region (see https://www.who.int/countries/), bcgindex (see http://www.bcgatlas.org/), pop65 (see https://data.worldbank.org/indicator/SP.POP.65) and lntb10 (see http://www.bcgatlas.org/). The whole data set is also submitted to the journal’s repository. In this place, we would like to mention that the most of the countries with low income levels (annual per capita income less than $825 USD) reported zero deaths attributed to COVID-19 till May 30, 2020, and for this reason, specific 104 countries are selected for data analysis, and the data with incomplete observations is not considered As indicated in Section 1, we here formulate a regression equation with the mortality due to COVID-19 as the response variable and motivated by the study of Takahashi (2020), we consider the IVs for the following three covariates :

**bcgindex:** The number of years a country has included BCG vaccine in its national immunization program. 1: All individuals received mandatory vaccinations; 0 : BCG neither previously nor currently mandatory; Values between 0 and 1: BCG previously mandatory but now discontinued. See Data Appendix for details of the construction of the BCG index. The definition is taken from Takahashi (2020).
**region:** The World Health Organization (WHO) regional classification was used here. 1: African Region; 2: South-East Asia Region; 3: East-Mediterranean Sea Region; 4: Western-Pacific Asia Region; 5: Region of America; 6: European Region. The definition is taken from Takahashi (2020).
**pop65:** The ratio of the population over 65 years of age. The definition is taken from Takahashi (2020).

Another covariate is **lntb10**, which is defined as the number of TB infections per 100,000 people in logarithmic scale, i.e., this variable is approximately the same as the **LTBI**.

As Takahashi (2020) did, we here test the hypothesis if LTBI is associated with reduced COVID-19 mortality and present the data analysis using IV estimation for the modelling of COVID-19 data based on the aforesaid four covariates, where **bcgindex**, **region** and **pop65** are considered as IVs, and **lntb10** is considered as original explanatory variable. We extend it further and use two types of IVs to demonstrate how the GIVE statistic can help in deciding the fitting of model and choice of IVs for a better fit. Here the IVs following Wald’s (*Z*_1_) and Durbin’s (*Z*_2_) methods are generated for all the explanatory variables in both the models. The following values of GIVE statistics are obtained:

For the multiple linear regression model with `“bcgindex”`, `“region”` as IVs, `“lntb10”` as the original variable, and the mortality due to COVID-19 as the response variable, we obtain the values of the GIVE statistics as 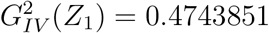 and 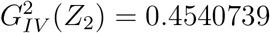. Moreover, using the *Z*_1_ type IV, the estimate of the coefficient associated with `“lntb10”` is −2.887569, and using the *Z*_2_ type IV, the estimate of the coefficient associated with `“lntb10”` is −2.7774120. The negative sign of the coefficient of `“lntb10”` and the relatively large values of 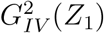 and 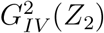 indicate that the LTBI may really protect against mortality due to COVID-19.

Next, for the multiple linear regression model with `“bcgindex”, “region”` and `“pop65”` as IVs, `“lntb10”` as the original variable, and the mortality due to COVID-19 as the response variable, we obtain the values of the GIVE statistics as 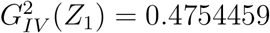 and 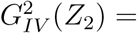 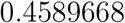. Moreover, using the *Z*_1_ type IV, the estimate of the coefficient associated with `“lntb10”` is −2.887569, and using the *Z*_2_ type IV, the estimate of the coefficient associated with `“lntb10”` is −0.03867306. Again, the negative sign of the coefficient of `“lntb10”`, and the larger values of 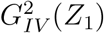 and 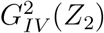 indicate that the LTBI may strongly protect against COVID-19 mortality for the population with age more than 65.

Overall, it clearly indicates that the multiple linear regression fits the COVID-19 data well, and the LTBI has really an impact on the reduction of mortality due to COVID-19; in particular, for the elder people, who are 65 years old or more.

## 6 Recommendations for using GIVE statistics

Based on the results in subsections 4.1, 4.2 and Section 5, the following recommendations are being made.

- The proposed GIVE statistics are capable of measuring the goodness of fitted models in case the regression coefficients are estimated by IV estimation method.
- The proposed GIVE statistic works well in the models with as well as without an intercept term.
- The performance of GIVE statistics depends upon the choice of IV, sample size and variance of random errors.
- The proposed GIVE statistics are capable of judging the the appropriate choice of IVs in order to have a better fitted model.
- The proposed GIVE statistics are capable of judging the variable selection, i.e., if the explanatory variables added in the model are relevant or not.
- The values of GIVE statistics can be used in any biological, biostatistical or econometric models, where IV is used.
- The use of GIVE statistic is not recommended when either the variance of random errors (or variance of the measurement error) is large or sample size is too low.
- The GIVE statistic performs better when the variance of random errors (or the variance of the measurement error) is not too large and/or sample size is reasonably large, but it also depends upon the choice of IVs.
- The GIVE statistics can be used in any real data application.

## 7 Supplementary Material

The supplementary file of this article contains the proofs of Theorem 1, Theorem 2, Theorem 3, the results of simulation study for multiple linear regression model and measurement error model.

## Acknowledgement

The authors are partially supported by their MATRICS grant of SERB (Files no : MTR/2019/000039 and MTR/2019/000033, respectively), Government of India.

## References

[1] Anderson, T.W. (2003): An Introduction to Multivariate Statistical Analysis, John Wiley.

[2] Baiocchi, M., Cheng, J. and Small, D.S. (2014) Tutorial in biostatistics: instrumental variable methods for causal inference. Statistics in Medicine, 33, 2297–2340.

[3] Bloom, N., Bond, S. and Van Reenen, J. (2007) Uncertainty and Investment Dynamics. The Review of Economic Studies, 74, 391–415.

[4] Bowden, R. L. and Turkington, D. A. (1984) Instrumental Variables, Cambridge University Press, Melbourne.

[5] Burgess, S., Small, D. S., Thompson, S. G. (2017) A review of instrumental variable estimators for Mendelian randomization. Statistical Methods and Medical Research, 26, 2333–2355.

[6] Cheng, C. L. and Van Ness, J. W. (1999) Statistical Regression with Measurement Error. London: Arnold and New York: Oxford University Press.

[7] Dai, J. Y., Chan, K. C. and Hsu, L. (2014) Testing concordance of instrumental variable effects in generalized linear models with application to Mendelian randomization. Statistics in Medicine, 33, 3986–4007.

[8] Didelez, V. and Sheehan, N. (2007) Mendelian randomization as an instrumental variable approach to causal inference. Statistical Methods and Medical Research, 16, 309–330.

[9] Eric, J. Tchetgen, Stefan Walter, Stijn Vansteelandt, Torben Martinussen and Maria Glymour (2015) Instrumental variable estimation in a survival context. Epidemiology, 26, 402–410.

[10] Fuller, W. A. (1987) Measurement Error Models. New York: Wiley,

[11] Li, J., Fine, J. and Brookhart, A. (2015) Instrumental variable additive hazards models. Biometrics, 71, 122–130.

[12] Martinussen, T. and Vansteelandt, S. (2020) Instrumental variables estimation with competing risk data. Biostatistics, 21, 158–171.

[13] Pesaran, M. H. and Smith, R. J. (1992) A Generalized *R*^2^ Criterion for Regression Models Estimated by the Instrumental Variables Method. Econometrica, 62, 705–710.

[14] Rao, C. R., Toutenburg, H., Shalabh and Heumann, C. (2008): Linear Models and Generalizations, Least Squares and Alternatives, 3rd edition, Springer, Berlin, Heidelberg.

[15] Soliman, R., Brassey, J., Pluddemann, A. and Henegahn, C. (2020): Does BCG vaccination protect against acute respiratory infections and COVID-19? A rapid review of current evidence. CEBM Working Paper 23 April 2020, https://www.cebm/wp-content/uploads/2010/04/BCG.jpg; 2020.

[16] Takahashi, H. (2020) Role of latent tuberculosis infections in reduced COVID-19 mortality: Evidence from an instrumental variable method analysis. Medical Hypotheses, 144, 110214.

[17] Tchetgen, E. J. (2014) A note on the control function approach with an instrumental variable and a binary outcome. Epidemiol Methods, 3, 107–112.

[18] Uddin, M. J., Groenwold, R. H., Ton de Boer, Belitser, S. V., Roes, K. C. and Klungel, O. H. (2015) Instrumental Variable Analysis in Epidemiologic Studies: An Overview of the Estimation Methods. Pharmaceutica Analytica Acta, 6, 1000353.

[19] Wansbeek, T. and Meijer, E. (2000) Measurement Error and Latent Variables in Econometrics, Elsevier Science, Amsterdam.

[20] Windmeijer, F. (1995) A note *R*^2^ in the instrumental variables model. Journal of Quantitative Economics, 11, 257–261.

